# DriveWays: A Method for Identifying Possibly Overlapping Driver Pathways in Cancer

**DOI:** 10.1101/2020.04.01.015388

**Authors:** Ilyes Baali, Cesim Erten, Hilal Kazan

## Abstract

**Motivation:** The majority of the previous methods for identifying cancer driver modules output non-overlapping modules. This assumption is biologically inaccurate as genes can participate in multiple molecular pathways. This is particularly true for cancer-associated genes as many of them are network hubs connecting functionally distinct set of genes. It is important to provide combinatorial optimization problem definitions modeling this biological phenomenon and to suggest efficient algorithms for its solution.

**Results:** We provide a formal definition of the Overlapping Driver Module Identification in Cancer (ODMIC) problem. We show that the problem is NP-hard. We propose a seed-and-extend based heuristic named DriveWays that identifies overlapping cancer driver modules from the graph built from the IntAct PPI network. DriveWays incorporates mutual exclusivity, coverage, and the network connectivity information of the genes.

We show that DriveWays outperforms the state-of-the-art methods in recovering well-known cancer driver genes performed on TCGA pan-cancer data. Additionally, DriveWays’s output modules show a stronger enrichment for the reference pathways in almost all cases. Overall, we show that enabling modules to overlap improves the recovery of functional pathways filtered with known cancer drivers, which essentially constitute the reference set of cancer-related pathways.

**Availability:** The data, the source code, and useful scripts are available at: https://github.com/abu-compbio/DriveWays

**Supplementary information:** Supplementary data are available at *Biorxiv*.

## 1 Introduction

Recent advances in high-throughput DNA sequencing technology have allowed several projects such as The Cancer Genome Atlas (TCGA) to systematically generate genomic data for thousands of tumors across many cancer types [1]. A key fundamental challenge in cancer genomics is to distinguish functional mutations that drive tumorigenesis, or *drivers*, from the numerous non-functional *passenger* mutations that occur randomly but that are not important for cancer development. Such a challenge is further complicated by the highly interactive nature of genes/proteins, thus necessitating the identification not only of such drivers but also of modules consisting of webs of drivers culpable in cancer initation and progression. Several computational approaches have been proposed for the cancer driver module identification problem and they can be categorized according to the types of biological data they utilize and the proposed optimization functions to model the underlying biological problem.

Early approaches for driver module analysis have primarily utilized the mutation data, in particular the frequency of mutations [2, 3, 4] or the positional clustering of mutations [5]. These methods can provide limited results as cancer genomes exhibit extensive mutational heterogeneity. Multiple approaches have been proposed to alleviate this problem. Rather than using mutation frequencies directly, Hotnet2 applies a random walk strategy to diffuse the mutation frequencies throughout the network and then identifies driver modules as strongly connected components of the resulting network [6]. Another direction is to utilize the concept of mutual exclusivity, the fact that multiple alterations in the same functional pathway occur less frequently because of diminished selective pressure. There exist methods that calculate all pairwise mutual exclusion scores [7, 8]. However, most methods limit the search space by using prior interaction knowledge. For instance, Ciriello *et al.* test each clique in the interaction network against random permutations to estimate the significance of mutation overlaps [9]. Vandin *et al.* propose a score that rewards coverage and penalizes mutation overlaps, and then searches for a set of genes that maximizes this score [10]. The same scoring function is also utilized by follow-up methods with an extension on the search technique [11, 12]. Babur *et al.* improves over the scoring function of [10] by fixing the bias towards highly altered genes [13]. Their proposed statistical mutual exclusivity test is used within a greedy search to identify groups of genes with high mutual exclusivity. MEMCover is also based on a greedy iterative seed-and-extend heuristic where a function that integrates coverage, mutual exclusivity and confidence values of interactions in the network is maximized [14]. MEXCOWalk extends Hotnet2’s random walk strategy by introducing edge weights that include mutual exclusity and coverage [15].

A common theme in almost all the mentioned cancer module identification methods is to search for nonoverlapping modules. However, biological pathways often overlap since proteins may carry out more than one function or belong to more than one protein complex [16]. Protein multifunctionality can also be considered as a means to coordinate multiple cellular activities serving as switches between pathways. As such, current methods that ultimately aim to provide a subset of existing biological pathways that are associated with cancer assume a problem definition that does not reflect the nature of biological pathways. To the best of our knowledge, only two previous methods provide possibly overlapping cancer driver modules, MEMCover [14] and ModulOmics [17]. In the former, no criteria for overlaps is included in the main search procedure which produces nonoverlapping output modules. The possible overlaps are only achieved via an optional post-processing step and no performance evaluations are done for this setting. ModulOmics integrates PPI network proximity, mutual exclusivity of DNA alterations, and RNA level coregulation and coexpression, into a single probabilistic framework, by simultaneously optimizing over all four model components. A significant shortcoming of ModulOmics is the lack of control over the amount of overlaps between the driver modules. For instance, for breast cancer, ModulOmics provides its top 50 modules, ranging in size 2 to 4, in the published results. Among these top 50 modules many of them are almost the same; 402 pairs differ only by one gene.

On the other hand there are some related problems in a wide range of areas including biological networks and social networks, such as protein complex identification or community detection, where overlapping module identification is an important research topic; see [18] for a survey on the topic. Shih *et al.* propose a soft variation of regularized Markov clustering to enable the identification of overlapping clusters in PPI networks [19]. ClusterOne uses a modularity metric in a weighted graph to guide the search for finding possibly overlapping subgraphs that correspond to protein complexes [20]. Bennett *et al.* propose a mixed integer nonlinear programming model to transform non-overlapping modules to overlapping, and apply this method to PPI networks of multiple organisms [21]. Although the proposed methods provide valuable insight on overlapping module constructions in the general setting, they are not designed for finding disease-associated modules. Modeling disease association requires extensive changes both in the input data and on the search procedure.

We propose DriveWays designed to identify potentially overlapping cancer driver modules. DriveWays uses a seed-and-extend strategy on a PPI network where it adds or removes gene sets based on a novel scoring function that includes coverage and mutual exclusivity of the module. The sizes of output modules can be controlled via appropriate parameters. We show that DriveWays improves over existing methods in the recovery of known cancer genes and more importantly in the recovery of pathways of known cancer driver genes. For the latter, we propose novel evaluation strategies that should prove useful for further research in this area.

## 2 Methods

Given the mutations data from a cancer cohort and a H. Sapiens PPI network, the informal goal is to extract from the PPI network subsets of genes (modules) that best reflect pathways related to the cancer under study. Ideally, these should correspond to the important causal functional pathways of driver genes of the relevant cancer. We first provide a computational problem definition to model this biological phenomenon. We discuss the computational complexity of the problem and provide an efficient greedy heuristic algorithm.

### 2.1 Problem Definition

Let *G* = (*V, E*) represent the PPI network where each vertex *u*_*i*_ ∈ *V* denotes a gene *g*_*i*_ whose expression gives rise to the corresponding protein in the network and each undirected edge (*u*_*i*_, *u*_*j*_) ∈ *E* denotes the interaction among the proteins corresponding to genes *g*_*i*_, *g*_*j*_. Henceforth assume *g*_*i*_ denotes both the gene and the corresponding vertex *u*_*i*_ in *G*. Let *S*_*i*_ denote the set of samples for which *g*_*i*_ is mutated and *S* denote the list of all such sets. Let *M* ⊆ *V* be a set of genes denoting a *module*. Let *G*(*M*) denote the subgraph of *G* induced by the vertices corresponding to genes in *M*. Since a driver pathway tends to be perturbed in a relatively large number of patients, one of the desired properties of each module is large *coverage* [14, 15, 6]. We define the coverage of *M* as, 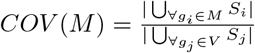. Several cancer driver module identification methods have additionally made use of the concept of *mutual exclusivity* [13, 14, 15, 9, 22]. It refers to the phenomenon that for a group of genes which exhibit evidence of shared functional pathway, simultaneous mutations in the same patients are less frequent than is expected by chance [7]. Formally, we define the mutual exclusivity of a module *M* as, 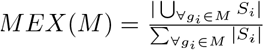. Combining the two functions, we define the *module score* of *M* as, *MS*(*M*) = *COV* (*M*) × *MEX*(*M*). Finally, for a set *D* of modules we define the *overlapping driver module set score* as, ODMSS(D) = Σ_∀*M*∈*D*_ ^*MS*(*M*)^.

Given as input a 4-tuple ≺ *G, S, δ*_*m*_, *δ*_*s*_ ≻, where *δ*_*m*_ and *δ*_*s*_ are integers, we define the *overlapping driver module identification in cancer (ODMIC)* problem as that of finding a set *D* of *possibly* overlapping modules that maximizes the *ODMSS*(*D*) and that satisfies the following:

- *Connectivity:* For each *M* ∈ *D*, *G*(*M*) is connected.
- *Uniqueness:* For each *M*_*i*_, *M*_*j*_ ∈ *D*, *M*_*i*_ ≠ *M*_*j*_.
- *Minimum Size: min*_∀*M*∈*D*_|*M*| = *δ*_*m*_.
- *Total Size:* Σ_∀*M*∈*D*_|*M*| = *δ*_*s*_.

We note that this definition is in part inspired by the cancer driver module identification problem definition of MEXCOWalk [15]. One crucial difference is that the MEXCOWalk definition does not allow overlaps. Secondly, due to the lack of overlaps, MEXCOWalk optimization score requires size normalizations in the contributions of MEX and COV. Furthermore the ODMIC scoring function is the sum of independent scores of modules and thus allows quite a different solution structure than that of MEXCOWalk. Finally, the size constraint of the output set of modules is with respect to the size of the set of unique genes in MEXCOWalk, whereas our problem definition applies the *Total Size* constraint which is determined by the sum of the sizes of the output modules. Such a choice allows flexible overlaps to be realized in an optimum solution. For instance, for *δ*_*s*_ = 10, a single module of size 10, two nonoverlapping modules of size 5 each, or two modules of size 5 with 4 common genes, all constitute legal instances in the solution space. Regardless of the differences in the problem definitions, we show that a reduction similar to the one employed in MEXCOWalk applies to this problem as well and that the problem in its generality is computationally intractable.

#### Theorem 2.1.

*The ODMIC problem is NP-hard.*

*Proof*. See the Supplementary Document.

The following lemma provides further intuition on the ODMIC problem by stating a fact regarding the structure of an optimum solution.

#### Lemma 2.2.

*There is an optimum solution D of the ODMIC problem on input instance* ≺ *G, S, δ*_*m*_, *δ*_*s*_ ≻, *where* |*M*| < 2*δ*_*m*_, ∀*M* ∈ *D*.

*Proof*. Let *D* be an optimum solution. We show that any *M* ∈ *D* with |*M*| ≥ 2*δ*_*m*_ can be split into two smaller modules *M*_1_, *M*_2_, each satisfying the *δ*_*m*_ constraint, such that *MS*(*M*_1_) + *MS*(*M*_2_) ≥ *MS*(*M*). Let *u*_1_, *s*_1_ denote respectively 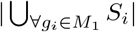 and 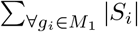. Let *u*_2_; *s*_2_ denote analogous values for *M*_2_. Since 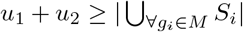 to show that 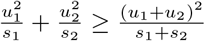, which holds trivially since (*u*_1_*s*_2_ − *u*_2_*s*_1_)^2^ ≥ 0.

Due to this structural property the ODMIC problem admits a pseudo-polynomial time algorithm under a certain setting.

#### Theorem 2.3.

*The ODMIC problem is solvable in pseudo-polynomial time for constant δ*_*m*_.

*Proof.* We propose a solution based on dynamic programming. Let *D* be an optimum solution of a given ODMIC input instance. By the *Minimum Size* constraint of the problem definition and by Lemma 2.2, we have *δ*_*m*_ ≤ |*M*| < 2*δ*_*m*_, for *M ∈ D*. Given a graph of *n* vertices, there are 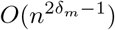 induced connected subgraphs with the allowed sizes. Since *δ*_*m*_ is constant, in an enumeration *M*_1_, *M*_2_ …, *M*_*p*_ of all such subgraphs we have *p* = *O*(*n*^*k*^), for constant *k*. Consider an optimum score table *c*, where *c*[*i, j*] indicates the optimum ODMSS score of an input instance consisting of subgraphs *M*_1_, *M*_2_, … *M*_*i*_ and the *Total Size* constraint set to *j*. Then *c*[*i, j*] = *max*(*c*[*i* − 1, *j*], *c*[*i* − 1, *j* − |*Mi*|] + *MS*(*M*_*i*_)). Thus the optimum solution can be found in time *O*(*n*^*k*^ × *δ*_*s*_).

Although the above result is valuable in providing a theoretical intuition regarding the solution structure, it is not fit for many practical settings. Efficient algorithms that may be suboptimal but that provide solutions close to optimum by making careful design choices with respect to the optimization criteria of the ODMIC problem are necessary.

### 2.2 DriveWays Algorithm

We provide a polynomial-time heuristic algorithm, DriveWays, for the ODMIC problem. It is based on a greedy seed-and-extend procedure on the input PPI network, that incorporates mutual exclusivity and coverage information. The pseudocode is provided in Algorithm 1. There are two main steps of the algorithm: (i) rank the genes with respect to the *MS* scores within the immediate neighborhoods (ii) initialize the module with the highest ranked seed and iteratively modify it by adding or removing a set of genes. The second step is repeated multiple times until we satisfy the ODMIC problem definition constraint regarding *Total Size*. Details are described in the following subsections.

### 2.2.1 Ranking the Seeds

Prioritizing cancer genes based on a combined score of coverage and mutual exclusivity has been employed in several previous approaches [15, 10, 14]. In line with our ODMIC problem definition, we similarly make use of coverage and mutual exclusivity values in the form of our module score definition. We first filter out the genes that are mutated in less than 1 % of the cohort. Then the remaining genes are sorted in nonincreasing order with respect to the module scores of their extended neighborhoods, that is, *MS*(*N*_*e*_(*g*)), where *N*_*e*_(*g*) denotes the set of neighbors of gene *g* in *G*, together with *g*. Such a score is basically a measure of how fit a gene is for further immediate growth with the neighbors. We note that we assessed the importance of our seed ranking procedure by rerunning DriveWays with randomly selected seed lists. We observe that the modules obtained with randomly selected seeds perform significantly worse than our original set of output modules, in terms of all the evaluation criteria considered in this study; see Supplementary Figures S2 and S3 for details.

### 2.2.2 Constructing Set of Driver Modules

We construct the set *D* of possibly overlapping modules through a greedy iterative module update procedure. For constructing a new module *M* to be added to *D*, we initialize *M* with the highest ranking seed that does not appear in an already existing module. We update *M* by either adding or removing certain gene(s) iteratively until such modifications no longer provide a gain to the current module in terms of the *MS* score. To check whether any gene additions to the current module provide a gain, we first construct a candidate set *CS*(*M*), from which the genes to be possibly added to *M* are selected. Let *N* (*g*_*i*_) denote the neighborhood of *g*_*i*_ in *G* and let 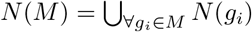. A gene *g*_*i*_ ∈ *N*(*M*) is added to *CS*(*M*), if it satisfies the following two conditions:

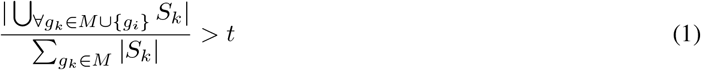

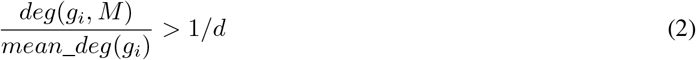

Inequality 1 relates the coverages of *M* with or without *g*_*i*_ to the mutual exclusivity of *M*. More specifically, it requires that the new coverage of the module with *g*_*i*_ should at least be a constant multiple *t* of the ratio of the old coverage to the old mutual exclusivity. In Inequality 2, *deg*(*g*_*i*_, *M*) denotes the degree of *g*_*i*_ in the subgraph of *G* induced by *M* ⋃ {*g*_*i*_}. On the other hand *mean_deg*(*g*_*i*_) is the average across *deg*(*g*_*i*_, *M*_*q*_) values, where *g*_*i*_ ∈ *M*_*q*_, for already existing *M*_*q*_ ∈ *D*. Thus by Inequality 2 a gene *g*_*i*_ is a candidate to be possibly added to the current module *M*, if it is well-connected to *M*, as compared to its connectivity to the already existing modules. Let *M*_*a*_ be the union of *M* with the set of genes *g*_*a*_ ∈ *CS*(*M*) that maximize *MS*(*M* ⋃ {*g*_*a*_}). Let *M*_*r*_ be the difference of *M* with the set of genes *g*_*r*_ ∈ *M* that maximize *MS*(*M* \ *g*_*r*_). The modules *M*_*a*_, *M*_*r*_ compete in *MS* improvement; if at least one improves over *MS*(*M*), the one with larger improvement is committed on *M*. If no improvement is achieved, the modifications of *M* are finalized and it is added into *D*. This procedure of module updates from a single seed are continued until the sum of the sizes of the modules in *D* reaches *δ*_*s*_.

**Algorithm 1.**
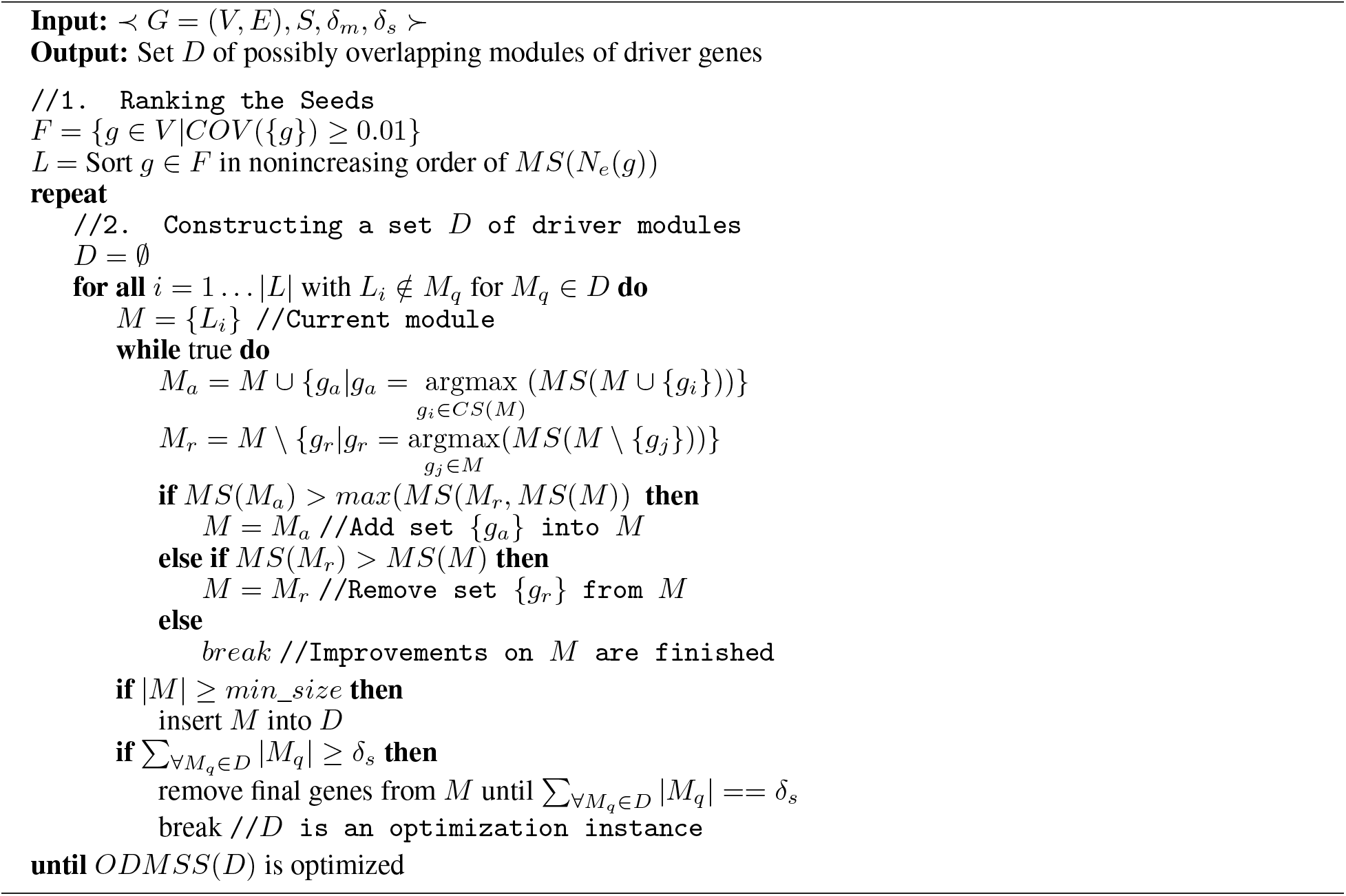
*DriveWays*

### 2.2.3 Optimizing with Respect to Parameters *t, d*

The constant multiple *t* in Inequality 1 is a parameter that indirectly controls the sizes of the output modules. Note that in the algorithm we do not explicitly control the module sizes in accordance with Lemma 2.2, since *t* achieves the same goal with more flexibility. On the other hand, the parameter *d* of Inequality 2 indirectly controls the overlap rate of the modules. An important feature of the algorithm is to set the parameters *t, d* automatically via an optimization function which chooses the instance of *D* that maximizes the main optimization function, *ODMSS*(*D*), among several instances produced through different *t, d* settings. The *repeat* loop of the main algorithm corresponds to this procedure implemented with the Bayesian Optimization (BayesOpt) procedure of [23].

## 3 Discussion of Results

We implemented the DriveWays algorithm in Python. The source code, useful scripts for evaluations, and all the input data are freely available as part of the supplementary material. We compare the results of DriveWays against those of four alternative methods. Among these alternatives three of them are knowledge-based cancer driver module identification methods: Hotnet2, MEMCover, and MEXCOwalk. As the fourth alternative method we employ ClusterOne, which is a representative algorithm for community detection in networks and outputs overlapping modules without any reference to cancer-related data. MEMCover is chosen due to its close connection to our work. It also optimizes mutual exclusivity and coverage of modules with a greedy seed-and-extend heuristic. Moreover, MEMCover is able to provide overlapping modules via a post-processing step. Hotnet2 is a good representative of heat diffusion based module finding algorithms though it only considers coverage, whereas MEXCOwalk improves over Hotnet2 by introducing edge weights that consider both mutual exclusivity and coverage.

### 3.1 Input data

All the methods except ClusterOne use the same input in the form of mutation data of available samples from TCGA and a PPI network. ClusterOne only uses the network information. We download the somatic aberration data from TCGA pan-cancer cohort preprocessed by [6]. The preprocessing procedure removes hypermutated samples and the genes with low expression throughout the tumor types. After this preprocessing, the dataset contains somatic aberrations for 11,565 genes in 3,110 samples. We perform two types of evaluations; one with all pan-cancer samples and one with breast cancer samples only. For the PPI network, we use the IntAct network downloaded from https://www.ebi.ac.uk/intact/ on Feb 11, 2019. The interactions with a confidence value less than 0.35 are filtered out. The final network contains 8,684 genes and 83,124 edges.

For evaluation purposes, three curated reference pathway sets are used: KEGG [24], Reactome [25], and BioCarta [26]. To obtain only significant cancer related pathways, we filter the reference pathways to include only *known cancer genes* and then we remove any resulting pathway with size less than *δ*_*m*_. Hereafter we denote each such set of reference pathways as *X*_*Y*_, where *X* is the employed pathway database and *Y* is the database of *known cancer genes* employed for filtering *X*. To compile known cancer genes for such filterings in pan-cancer evaluations, we use the COSMIC Cancer Gene Census (*CGC*) database [27]. However the *CGC* list lacks a complete annotation of cancer type information. As such, for the breast cancer evaluations, we employ the CancerMine database to construct the corresponding set [28]. CancerMine employs text-mining to catalogue cancer-associated genes through which it also extracts information about cancer types. We compile the list of CancerMine’s breast cancer-associated genes that have at least 3 citations and call it *CM3*.

Finally, for the GO term analysis of this section, we employ the go-basic.obo file from http://geneontology.org on June 26, 2019. We restrict the gene annotations to level 5 of the GO hierarchy by ignoring the higher-level annotations and replacing the deeper-level category annotations with their ancestors at the restricted level. We call the resulting terms as the *standardized GO terms*.

### 3.2 Parameter settings

A parameter applied commonly to all the methods under consideration is *δ*_*m*_ which is set to 3, as this constitutes a nontrivial minimum module size compatible with the problem definition. For DriveWays we find the *t* and *d* setting that maximizes ODMSS. We utilize the BayesOpt procedure implemented in scikit-optimize package to find these values in a time efficient manner [23]. We use the version 0.7.4 with the following setting of the arguments: *n*_*calls*_ = 30 and *acq_func* = *EI*. We search for the optimal value of *t* in the range [0.8, 1.2] and the optimal value of *d* in the range [2, 5]. Selected values are available in Table S5. For ClusterOne, we set the penalty term *p* and the overlap score threshold *w* to their default values, 2 and 0.8, respectively. For Hotnet2, the recommended value of 0.4 is used for the restart probability. Regarding MEXCOWalk, the default values are used for the restart probability (*β* = 0.4) and the mutual exclusivity threshold (*θ* = 0.7). For MEMCover, as recommended in the original study, the mutual exclusivity scores are obtained from type-restricted permutation test with all pan-cancer samples. Coverage parameter *k* is set to its default value of 15. *f*(*θ*) is a parameter that indirectly controls the module sizes in MEMCover. It is chosen such that the number of modules with size < *δ*_*m*_ is minimized.

### 3.3 Evaluations Omitting Modularity

Before performing any evaluations with respect to the specific grouping of the output genes into modules, we simply check whether our method recovers more known drivers when each output set under comparison is considered be a single set consisting of the union of the genes in all the output modules provided by each method. To provide a fair comparison between the methods outputting overlapping modules (DriveWays, ClusterOne, and MEMCover) and those providing nonoverlapping modules (HotNet2 and MEXCOWalk), rather than fixing the *δ*_*s*_ parameter for different methods, we obtain the results by varying the total number of unique genes, named *unique_genes*, from 100 to 1000 in steps of size 100. To achieve this for ClusterOne, MEMCover, and DriveWays we take the top ranking modules until the number of unique genes is equal to the *unique_genes*. For Hotnet2 and MEXCOWalk, we choose an edge weight threshold value such that removal of edges below this threshold value results in strongly connected components with total size equal to *unique_genes*. For each method and for varying values of *unique_genes* = 100, 200, …, 1000, we compute the intersection of the output gene lists, that is the union of the output modules, with the reference database of known drivers to calculate true positive and false positive rates. Figure 1 plots the Receiver Operating Characteristic (ROC) curves obtained from each method for the pan-cancer data, taking *CGC* as the reference set. We observe that DriveWays performs better than all the other methods. Hotnet2 and ClusterOne perform considerably worse than the other methods. ClusterOne’s poor performance is expected since it does not employ any cancer-related information. The analogous plot for the breast cancer cohort with the *CM3* as the reference set can be found in Figure S5 of the Supplementary. Different from the pan-cancer result, MEXCOWalk slightly outperforms DriveWays and MEMCover which are tied as the second best performers.

**Figure 1:**
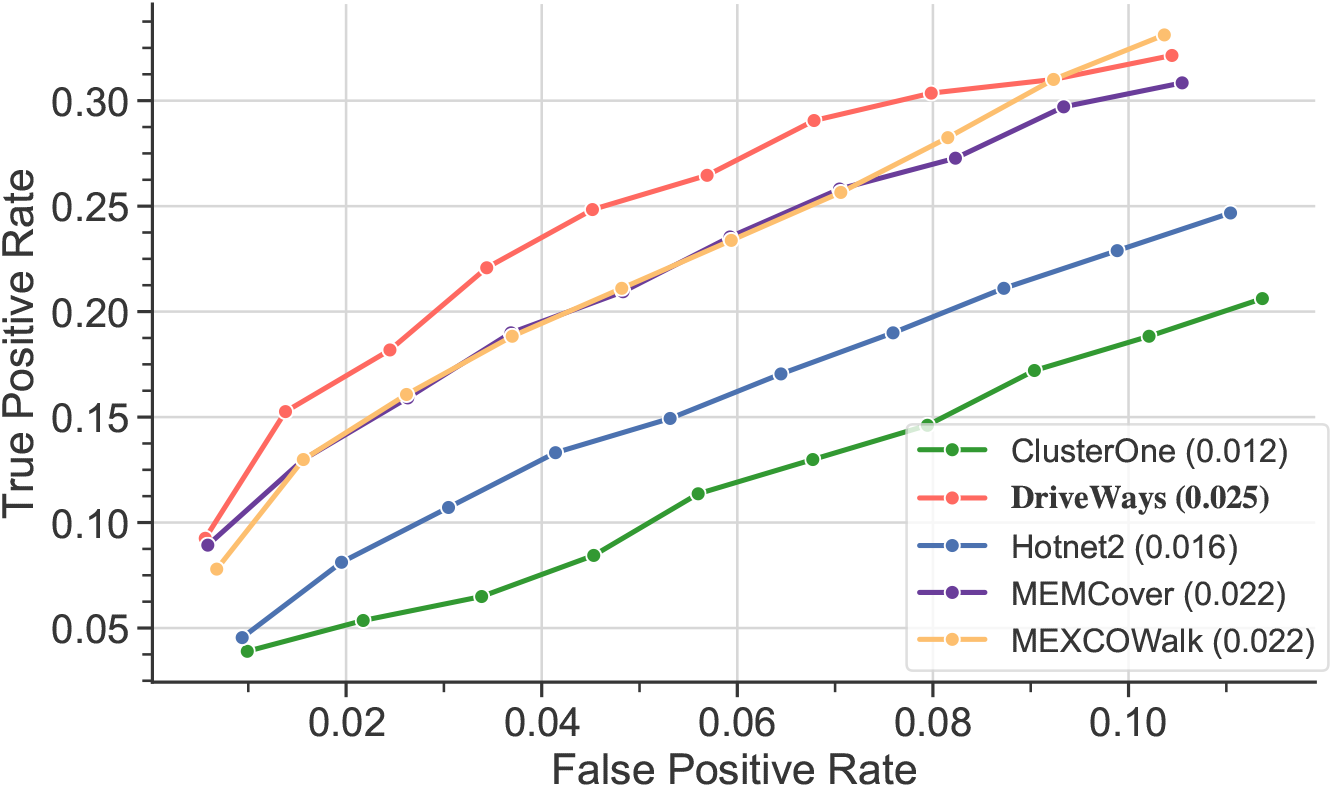
ROC curves calculated for *unique_genes* = 100, 200, …, 1000 from the output sets of modules of the methods under consideration.

### 3.4 Evaluations Based on Modularity

The goal of the cancer module identification methods is not only recovering the maximum number of known cancer drivers but more importantly providing them as groups of genes that share the same molecular functions or pathways. Since providing modules that cover all cancer-related pathways is critical, we set the main goal of the evaluation procedures as recovering each *set of reference pathways*, that is KEGG_*Y*_, Reactome_*Y*_, or Biocarta_*Y*_, where *Y* is *CGC* for the pan-cancer evaluations and *CM3* for the breast cancer evaluations. Therefore for the rest of the evaluations, *δ*_*s*_ parameter is set to Σ_∀*M*∈*D*′_|*M*|, where *D*′ indicates the corresponding set of reference pathways. This value is 1771 for KEGG_*CGC*_ and 845 genes for KEGG_*CM*3_; 3368 and 1416 genes for Reactome’s respective filtrations; 1173 and 626 genes for Biocarta’s respective filtrations. The desired outputs with the corresponding *δ*_*s*_ values for different methods can be achieved similar to the approach desribed in the previous subsection for the *unique_genes*. Note that upon setting *δ*_*s*_ to match the corresponding value from a specific set of reference pathways for all the methods, each method itself has the flexibility to choose how many unique genes it provides in its output, which in turn is correlated with the sizes of the output modules and the degree of overlaps among them.

### 3.4.1 Statistics on Output Modules and the Sets of Reference Pathways

The first statistic we provide is regarding the main optimization goal of our method, that is the overlapping driver module set score (ODMSS). Figure 2 shows that DriveWays predicted modules have significantly higher ODMSS values than the output modules of all the other methods. Additionally, the fairly large ODMSS scores observed for sets of reference pathways support the validity of the ODMSS as an objective function. Among the rest of the methods, ClusterOne’s performance is impressive considering that it is not a cancer-specific module identification method; it provides the fourth best performance surpassing Hotnet2. Rather than its module growth procedure, this performance could in part be due to ClusterOne’s seed ranking procedure which is based on the degree of the genes in the network. Using only this seed list as the output set of genes achieves a performance that is even better than that of ClusterOne itself in terms of the CGC overlap evaluations of the previous subsection that considers the union of output modules; see Supplementary Figure S4. Since most CGC genes also have high coverage scores, it is not surprising to observe that ClusterOne modules result in a high ODMSS value. For this metric, Hotnet2 performs the worst among the considered methods presumably due to its very large modules, as large modules are likely to show poor mutual exclusivitiy.

**Figure 2:**
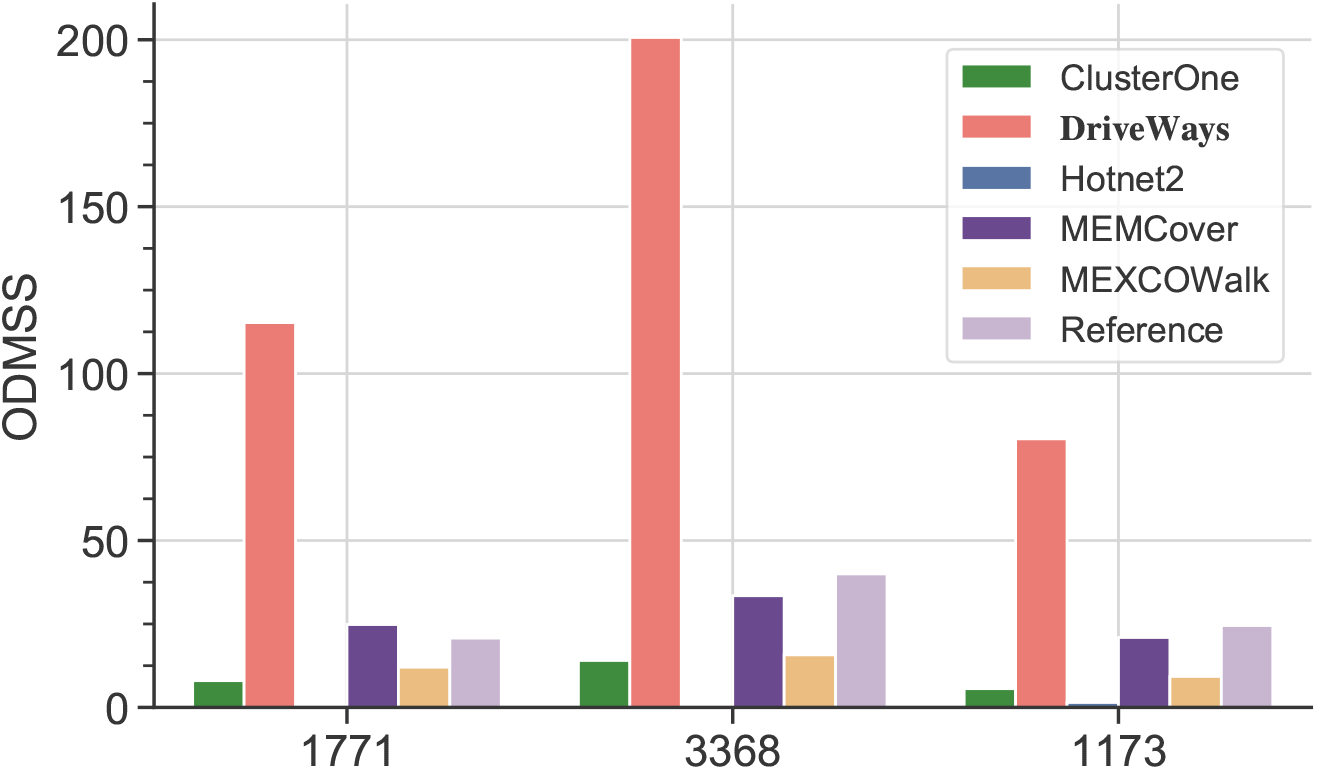
ODMSS values for the outputs of all the methods when *δ*_*s*_ is set to shown values.

We next provide certain statistics on the average sizes and the overlap rates of the output modules obtained for the methods under consideration. Figure 3-A shows the average module sizes when *δ*_*s*_ is set to KEGG_*CGC*_, Reactome_*CGC*_, and Biocarta_*CGC*_ sizes. The same plot also includes the average module sizes of the three sets of reference pathways themselves for comparison. We observe that Hotnet2 has the largest average module size which is significantly larger than those of the other methods. The second and the third largest average module sizes are obtained with ClusterOne and MEXCOWalk outputs, respectively. MEMCover and DriveWays have similar average module sizes that are smaller than the others. Further detailed statistics on the number of output modules and the number of unique genes in all the output modules pertaining both to the outputs of alternative methods and also to the actual sets of reference pathways themselves can be found in Tables S1 and S2 of the Supplementary.

**Figure 3:**
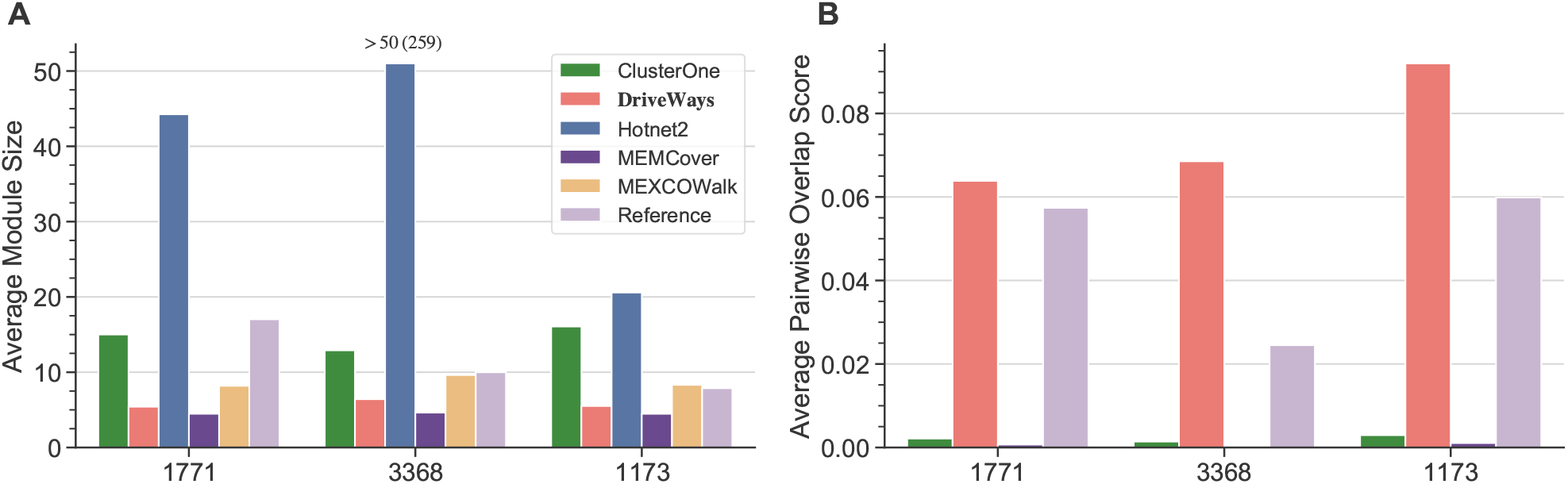
A) Average module sizes in the outputs of the methods under consideration for the shown *δ*_*s*_ values. B) Corresponding average pairwise overlap scores.

We quantify the degree of overlap among the output modules by calculating a *pairwise overlap score*, as previously defined in [29]. For two modules *M*_*i*_ and *M*_*j*_, the pairwise overlap score is calculated as, 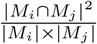. We calculate the of all such pairs. sum of the pairwise overlap scores for all pair of modules and normalize it by dividing by the number of all such pairs Figure 3-B shows the resulting average pairwise overlap scores of all the methods and the sets of reference pathways. Here, Hotnet2 and MEXCOWalk are excluded as they provide non-overlapping modules. We observe that modules of DriveWays overlap with each other more compared to the modules of other methods and that this overlap is quite similar to the overlap of the sets of reference pathways themselves.

Analogous statistics regarding the *δ*_*s*_ setting with respect to the KEGG_*CM*3_, Reactome_*CM*3_, and Biocarta_*CM*3_ pathway sizes and the corresponding ODMSS scores on the breast cancer data show quite similar results; see Supplementary Document for the relevant plots.

### 3.4.2 Recovering Sets of Cancer-associated Reference Pathways

A given set of predicted modules is evaluated by assessing how well they match and cover a set of reference pathways. Let {*M*_1_, …, *M*_*m*_}, {*R*_1_, … , *R*_*n*_ denote the set of predicted modules and the set of reference pathways, respectively. We introduce three measures to quantify the similarity between a predicted module *M*_*i*_ and a reference pathway *R*_*j*_.

#### Overlap score

We calculate the overlap score between a module and a pathway as the pairwise overlap score defined in the previous subsection, replacing *M*_*j*_ with the reference pathway *R*_*j*_ in the formula.

#### Hypergeometric test q-value

A hyper-geometric enrichment test is used to evaluate the significance of the intersection of *M*_*i*_ with *R*_*j*_. Adjusted p-values (also called q-values) are calculated with False Discovery Rate (FDR) correction [30].

#### GO consistency score

This score has been previously employed for the evaluation of PPI network alignment algorithms [31]. Let 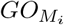 and 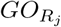 denote the union of the standardized GO terms obtained from the GO annotations of the genes in *M*_*i*_ and *R*_*j*_, respectively. GO consistency score of *M*_*i*_ and *R*_*j*_ is defined as, 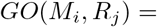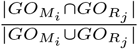.

Rather than identifying the enriched pathways for each module separately, we use an evaluation procedure which ensures that the set of predicted modules as a whole provides a good match to the whole set of reference pathways. To this end, the first metric we use is based on *Maximum Weighted Maximum Cardinality Matching (MWMCM)*. To identify MWMCM, we first create a bipartite graph containing nodes corresponding to the predicted modules on the one side and nodes corresponding to the reference pathways on the other side. The edge weights between a predicted module *M*_*i*_ and a reference module *R*_*j*_ are computed using one of the three similarity measures defined above. The overlap score and the GO consistency score of *M*_*i*_ and *R*_*j*_ can each be directly used as edge weights between the corresponding nodes of the bipartite graph. To use the q-values as edge weights, we transform them by taking the −*log*_10_ of the values so that larger ones correspond to better matches. Also, if the q-value is > 0.05, we instead assign zero as the edge weight as this corresponds to a non-significant match. Once the bipartite graph is formed, we find the MWMCM; that is, we find a subset of edges such that each predicted module and reference pathway is incident on at most one selected edge, the number of such selected edges is maximum (maximum cardinality matching), and the sum of the weights of selected edges is maximized among all maximum cardinality matchings. Lastly, we calculate the average weight of the edges in the resulting matching. We call this score Maximum Matching Ratio (MMR), as in [20]; see Figure S10 for a plot depicting how MMR is computed. We emphasize the fact that we employ a complete bipartite graph where zero-weight edges are also included, since excluding such edges could provide misleading results. For instance, consider a scenario where only one of the output modules is a perfect match to a reference pathway and the remaining modules show no similarity under any defined measure with any member of the set of reference pathways. When there is no similarity, the weights of edges connected to those modules would be zero. If zero-weight edges are removed the hypothetical method providing such an output would get a perfect MMR value of 1 even though only one of its predicted modules can be considered “good”; see Figure S11 for a toy example. On the other hand, calculation of MMR with zero-weight edges ensures that every predicted driver module is enriched for a functional pathway important for cancer.

Figure 4 displays the MMR results calculated with the three similarity measures. MEMCover has a slightly better MMR score than DriveWays under the GO consistency similarity measure when Reactome_*CGC*_ is used as the reference. In all the other evaluations, DriveWays gives higher MMR values than the competing methods. MEMCover ranks the second in most cases except for the MMR score under the q-value similarity measure when Reactome_*CGC*_ or Biocarta_*CGC*_ is used as the reference. In these two cases, ClusterOne performs better than MEMCover. Hotnet2 performs the worst in all evaluations. In particular, Hotnet2’s MMR scores under the q-value measure are close to zero presumably due to its large-sized modules.

**Figure 4:**
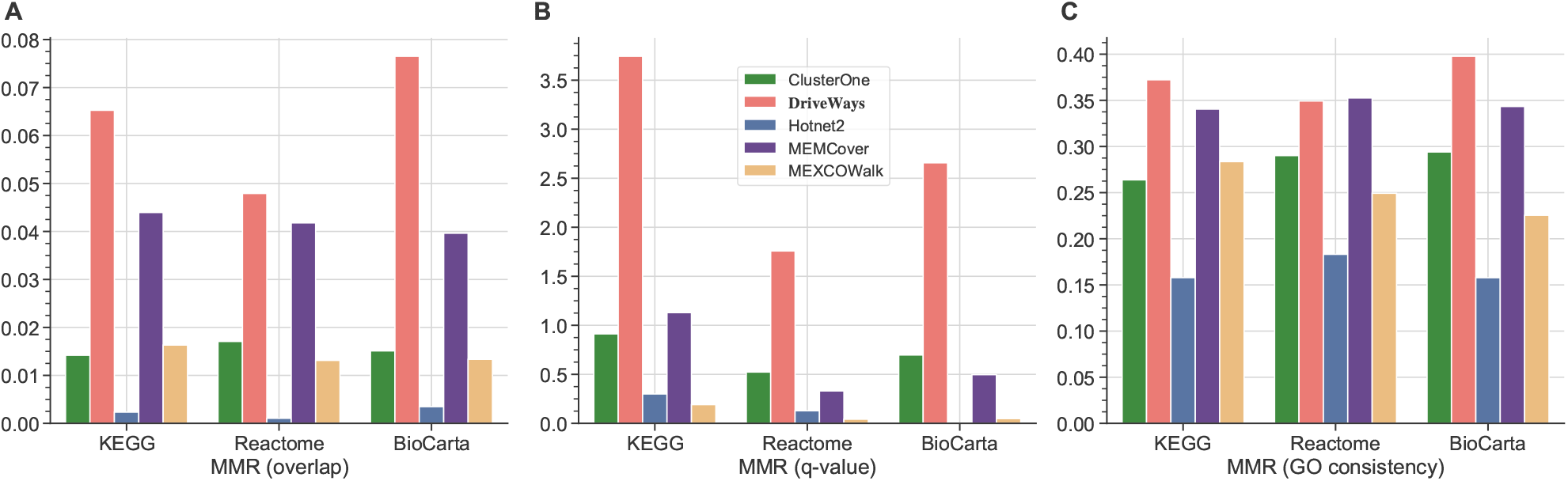
MMR scores of all methods calculated with three similarity metrics: A) Overlap score B) Hypergeometric test q-values C) GO consistency. The set of reference pathways at the x-coordinate of each plot correspond to KEGG_*CGC*_, Reactome_*CGC*_, and Biocarta_*CGC*_, from left to right.

Next, we utilize the precision and recall metrics to evaluate the predicted modules. A modified version of these metrics have been previously used in evaluating the quality of the predicted modules [32, 19]. To evaluate the precision of a method, for each of its predicted module *M*_*i*_, we find the *best match* in the set of reference pathways using one of the three similarity measures, overlap score, q-value, or GO consistency score. We evaluate recall similarly, but this time we find the *best match* of each reference pathway *R*_*j*_ among the predicted modules using one of the similarity measures. Identification of the best match for each predicted module and for each reference pathway is illustrated with a toy example in the Supplementary Figures S12 and S13. We plot the distribution of the best match scores across all the predicted modules and across all the reference pathways. Supplementary Figure S1 shows these distributions for each similarity metric, for each method, and for each set of reference pathways. We observe that the best match scores of DriveWays-predicted modules are significantly higher than the best match scores obtained with the other methods for all similarity metrics and for all sets of reference pathways. In terms of the best match scores of the set of reference pathways, DriveWays performs better in the majority of the cases with few exceptions. MEMCover performs slightly better than DriveWays under the overlap score similarity measure with respect to the Reactome_*CGC*_ set of reference pathways. Similarly, MEMCover performs better under the GO consistency similarity measure with respect to the Reactome_*CGC*_ and BioCarta_*CGC*_ sets of reference pathways. DriveWays’s slightly worse performance in terms of recall in these cases can be attributed to the relatively high pairwise overlaps of its output modules. Since *δ*_*s*_ is fixed, DriveWays outputs smaller number of unique genes as compared to the other methods and also as compared to the sets of reference pahtways. This is why some sets of reference pathways could have low best match scores since the genes in those reference pathways do not exist in the output modules of DriveWays. However, the performance difference between DriveWays and the rest of the methods in terms of precision dominates the performance difference between MEMCover and DriveWays in terms of recall. To illustrate this, we also compute an aggregate score by finding the average best match score across the predicted modules and the average best match score across the reference pathways. F1 scores obtained by the product of these two average values are shown in Supplementary Table S3. DriveWays has the highest F1 score in all the experiments illustrating its overall superiority with respect to precision and recall.

We repeat the same evaluations for the breast cancer data employing KEGG_*CM*3_, Reactome_*CM*3_, and Biocarta_*CM*3_ as the sets of reference pathways. Almost all the results regarding the pan-cancer data discussed here apply similarly in this setting as well; see the Supplementary Document for detailed plots.

### 3.4.3 Top DriveWays Modules Compared to Their Matches

We next investigate our top modules and their matched reference pathways more closely. We first look at our top 10 modules and their matched KEGG_*CGC*_ reference pathways in the context of MWMMC. First of all, we observe that all top 10 modules are incident on the set of edges selected for MWMMC. We further explore the reference pathways that are connected to the top 10 modules through these selected edges. Five of the ten such reference pathways directly correspond to a pathway of a specific cancer type: *Non small cell lung cancer, Bladder cancer, Glioma, Pancreatic cancer* and *Endometrial cancer*. Among the other matched reference pathways, *Cell cycle* pathway and the *p53 signalling pathway* are also strongly associated with cancer. We observe that the matches to *Cell cycle* and the *Non small cell lung cancer* pathways have the highest edge weights. For both matches, our predicted modules consist of five genes all of which also appear in the matched reference pathways. Another interesting match is observed between our seventh ranking module and the *Bladder cancer* pathway. Our predicted module contains six genes, four of which appear in the *Bladder cancer* pathway. VHL, a well known tumor-suppressor, is among the two genes that appear in our predicted module but not in the *Bladder cancer* pathway. Interestingly, among the cancer types in pan-cancer cohort, bladder cancer ranks second after renal cell carcinoma in terms of VHL’s mutation frequency.

We also explore the matches that are found within the context of precision. We explore the best matching reference pathways of top 20 predicted modules of DriveWays. Among the best matched KEGG_*CGC*_ pathways, we observe *Pathways in cancer* nine times, *Cell cycle* four times, *ERBB signaling pathway* three times, *p53 signalling pathway* twice, *MAPK signaling pathway* and *TGFB signaling pathway* once. Next, we identify the top 20 modules that contain a gene that do not appear in CGC reference. Since these genes are not in CGC, they also cannot appear in the corresponding best matched reference KEGG_*CGC*_ pathway. As such, we instead check whether these genes appear in the unfiltered versions of the corresponding KEGG pathways. We find four such genes that belong to four distinct predicted modules: LTBP1 appears in *TGFB signaling pathway*, SMC3 and SMC1A appear in *Cell cycle* pathway, and FLNA appears in *MAPK signaling pathway*. These genes could be novel cancer drivers as they appear in our top modules and they function in the same pathway as closely connected CGC genes in the network.

Lastly, in terms of recall, we identify the cancer related pathways in KEGG_*CGC*_ and find their best matches among the predicted modules of DriveWays and MEMCover; the two methods that can identify overlapping cancer driver pathways. When compared with MEMCover, except for the *Colorectal cancer pathway*, DriveWays’s predicted modules result in a better overlap score with cancer related KEGG_*CGC*_ pathways. Overall these results show that top DriveWays modules are enriched for cancer associated pathways in KEGG.

## 4 Conclusion

DriveWays is a novel method that incorporates network connectivity, mutual exclusivity, and coverage information to identify overlapping cancer driver modules. It does not require any additional parameters, other than the desired minimum size of a module and the sum of the sizes of all the modules, both of which should be intuitive properties for cancer biologists. Comparing against four state-of-the-art methods, we demonstrate the ability of DriveWays to identify modules enriched with known cancer genes, and also enriched for curated pathways containing only known cancer driver genes. Other key contributions of our work are both the introduction of an intuitive combinatorial optimization problem definition fairly representing the underlying biological phenomenon of identifying possibly overlapping cancer driver module identification and that of novel evaluation metrics suitable for evaluating methods proposed for solving it.

## Acknowledgements

The authors are listed in alphabetical order of their lastnames. We thank Rafsan Ahmed for his help in obtaining output modules of Hotnet2 and MEXCOWalk.

## References

[1] John N. Weinstein, Eric A. Collisson, Gordon B. Mills, Kenna R. Mills Shaw, Brad A. Ozenberger, Kyle Ellrott, Chris Sander, and et al. The cancer genome atlas pan-cancer analysis project. Nature Genetics, 45(10):1113–1120, 10 2013.

[2] C. Greenman, R. Wooster, and P. A. Futreal. Statistical analysis of pathogenicity of somatic mutations in cancer. Genetics, 173:2187–2198, 2006.

[3] G. Getz, H. Hofling, J. P. Mesirov, T. R. Golub, M. Meyerson, R. Tibshirani, and E. S. Lander. Comment on "The consensus coding sequences of human breast and colorectal cancers". Science, 317(5844):1500, Sep 2007.

[4] A. Youn and R. Simon. Identifying cancer driver genes in tumor genome sequencing studies. Bioinformatics, 27(2):175–181, Jan 2011.

[5] D. Tamborero, A. Gonzalez-Perez, and N. Lopez-Bigas. OncodriveCLUST: exploiting the positional clustering of somatic mutations to identify cancer genes. Bioinformatics, 29(18):2238–2244, Sep 2013.

[6] Mark D. M. Leiserson, Fabio Vandin, Hsin-Ta Wu, Jason R. Dobson, Jonathan V. Eldridge, Jacob L. Thomas, Alexandra Papoutsaki, Younhun Kim, Beifang Niu, Michael McLellan, Michael S. Lawrence, Abel Gonzalez-Perez, David Tamborero, Yuwei Cheng, Gregory A. Ryslik, Nuria Lopez-Bigas, Gad Getz, Li Ding, and Benjamin J. Raphael. Pan-cancer network analysis identifies combinations of rare somatic mutations across pathways and protein complexes. Nature Genetics, 47(2):106–114, 2015.

[7] Chen-Hsiang Yeang, Frank McCormick, and Arnold Levine. Combinatorial patterns of somatic gene mutations in cancer. The FASEB Journal, 22(8):2605–2622, 2008.

[8] C. A. Miller, S. H. Settle, E. P. Sulman, K. D. Aldape, and A. Milosavljevic. Discovering functional modules by identifying recurrent and mutually exclusive mutational patterns in tumors. BMC Med Genomics, 4:34, Apr 2011.

[9] Giovanni Ciriello, Ethan Cerami, Chris Sander, and Nikolaus Schultz. Mutual exclusivity analysis identifies oncogenic network modules. Genome Research, 22(2):398–406, February 2012.

[10] F. Vandin, E. Upfal, and B. J. Raphael. De novo discovery of mutated driver pathways in cancer. Genome Res., 22(2):375–385, Feb 2012.

[11] J. Zhao, S. Zhang, L. Y. Wu, and X. S. Zhang. Efficient methods for identifying mutated driver pathways in cancer. Bioinformatics, 28(22):2940–2947, Nov 2012.

[12] M. D. Leiserson, D. Blokh, R. Sharan, and B. J. Raphael. Simultaneous identification of multiple driver pathways in cancer. PLoS Comput. Biol., 9(5):e1003054, 2013.

[13] Özgün Babur, Mithat Gönen, Bülent Arman Aksoy, Nikolaus Schultz, Giovanni Ciriello, Chris Sander, and Emek Demir. Systematic identification of cancer driving signaling pathways based on mutual exclusivity of genomic alterations. Genome Biology, 16(1):45, Feb 2015.

[14] Yoo-Ah Kim, Dong-Yeon Cho, and T. M. Przytycka. MEMCover: integrated analysis of mutual exclusivity and functional network reveals dysregulated pathways across multiple cancer types. Bioinformatics, 31(12):i284–i292, 2015.

[15] Rafsan Ahmed, Ilyes Baali, Cesim Erten, Evis Hoxha, and Hilal Kazan. MEXCOwalk: mutual exclusion and coverage based random walk to identify cancer modules. Bioinformatics, 36(3):872–879, 08 2019.

[16] A. Zanzoni, D. M. Ribeiro, and C. Brun. Understanding protein multifunctionality: from short linear motifs to cellular functions. Cell. Mol. Life Sci., 76(22):4407–4412, Nov 2019.

[17] Dana Silverbush, Simona Cristea, Gali Yanovich-Arad, Tamar Geiger, Niko Beerenwinkel, and Roded Sharan. Simultaneous integration of multi-omics data improves the identification of cancer driver modules. Cell Systems, 8(5):456–466.e5, 2019.

[18] Jierui Xie, Stephen Kelley, and Boleslaw K. Szymanski. Overlapping community detection in networks: The state-of-the-art and comparative study. ACM Comput. Surv., 45(4):43:1–43:35, August 2013.

[19] Yu-Keng Shih and Srinivasan Parthasarathy. Identifying functional modules in interaction networks through overlapping Markov clustering. Bioinformatics, 28(18):i473–i479, 09 2012.

[20] Tamás Nepusz, Haiyuan Yu, and Alberto Paccanaro. Detecting overlapping protein complexes in protein-protein interaction networks. Nature Methods, 9(5):471–472, 2012.

[21] Laura Bennett, Aristotelis Kittas, Songsong Liu, Lazaros G Papageorgiou, and Sophia Tsoka. Community structure detection for overlapping modules through mathematical programming in protein interaction networks. PloS One, 9(11):e112821, 2014.

[22] Phuong Dao, Yoo-Ah Kim, Damian Wojtowicz, Sanna Madan, Roded Sharan, and Teresa M. Przytycka. Bewith: A between-within method to discover relationships between cancer modules via integrated analysis of mutual exclusivity, co-occurrence and functional interactions. PLoS Computational Biology, 13(10), 2017.

[23] Tim Head, MechCoder, Gilles Louppe, Iaroslav Shcherbatyi, fcharras, Zé Vinícius, cmmalone, Christopher Schröder, nel215, Nuno Campos, Todd Young, Stefano Cereda, Thomas Fan, Justus Schwabedal, Hvass-Labs, Mikhail Pak, SoManyUsernamesTaken, Fred Callaway, Loïc Estève, Lilian Besson, Peter M. Landwehr, Pavel Komarov, Mehdi Cherti, Kejia (KJ) Shi, Karlson Pfannschmidt, Fabian Linzberger, Christophe Cauet, Anna Gut, Andreas Mueller, and Alexander Fabisch. scikit-optimize/scikit-optimize: v0.5.1 - rerelease, February 2018.

[24] M. Kanehisa, S. Goto, S. Kawashima, and A. Nakaya. The KEGG databases at GenomeNet. Nucleic Acids Res., 30(1):42–46, Jan 2002.

[25] A. Fabregat, S. Jupe, L. Matthews, K. Sidiropoulos, M. Gillespie, P. Garapati, R. Haw, B. Jassal, F. Korninger, B. May, M. Milacic, C. D. Roca, K. Rothfels, C. Sevilla, V. Shamovsky, S. Shorser, T. Varusai, G. Viteri, J. Weiser, G. Wu, L. Stein, H. Hermjakob, and P. D’Eustachio. The Reactome Pathway Knowledgebase. Nucleic Acids Res., 46(D1):D649–D655, 01 2018.

[26] S. M. Wimalaratne, N. Juty, J. Kunze, G. Janee, J. A. McMurry, N. Beard, R. Jimenez, J. S. Grethe, H. Hermjakob, M. E. Martone, and T. Clark. Uniform resolution of compact identifiers for biomedical data. Sci Data, 5:180029, 05 2018.

[27] Zbyslaw Sondka, Sally Bamford, Charlotte G. Cole, Sari A. Ward, Ian Dunham, and Simon A. Forbes. The cosmic cancer gene census: describing genetic dysfunction across all human cancers. Nature Reviews Cancer, 18(11):696–705, 2018.

[28] Jake Lever, Eric Y. Zhao, Jasleen Grewal, Martin R. Jones, and Steven J. M. Jones. Cancermine: a literature-mined resource for drivers, oncogenes and tumor suppressors in cancer. Nature Methods, 16(6):505–507, 2019.

[29] G. D. Bader and C.W. Hogue. An automated method for finding molecular complexes in large protein interaction networks. BMC Bioinformatics, 4(2):10.1186/1471-2105-4-2, 2003.

[30] Y. Benjamini and Y Hochberg. Controlling the false discovery rate: a practical and powerful approach to multiple testing. J of Royal Stat Society, 57(1):289–300, 1995.

[31] Ferhat Alkan and Cesim Erten. BEAMS: backbone extraction and merge strategy for the global many-to-many alignment of multiple PPI networks. Bioinformatics, 30(4):531–539, 12 2013.

[32] Guimei Liu, Limsoon Wong, and Hon Nian Chua. Complex discovery from weighted PPI networks. Bioinformatics, 25(15):1891–1897, 05 2009.

