## Supplementary Material for "DriveWays: A Method for Identifying Possibly Overlapping Driver Pathways in Cancer"

#### 1.1 Proof of Theorem 1

**Theorem 1.1.** *Overlapping driver module identification in cancer is NP-hard.*

*Proof.* The transformation is from *Set Packing* which is NP-complete [Garey and Johnson(1979)Garey and Johnson]. In the Set Packing problem, given a collection $C$ of finite sets and a positive integer $K \leq |C|$, the problem is to find out whether $C$ contains at least $K$ mutually disjoint sets. The problem is NP-hard even when the size of each set is at most 3, which can easily be extended to the setting where the size of each set is exactly 3. Given an input to the Set Packing problem within this setting in the form of $K$ and $C$ such that for each $S \in C$, $|S| = 3$, we generate $G$ as a complete graph on $|C|$ vertices, corresponding to the such that each finite set in $C$ corresponds to a set of samples $S_i$ for which gene $g_i$ is mutated. We set both $\delta_s$ and $min\_module\_size$ to $K$. The answer to the Set Packing problem is Yes, if and only if the maximized score of the overlapping driver module identification in cancer problem is exactly $\frac{3 \times K}{|\bigcup_{\forall g_i \in V} S_i|}$.

Note that the same reduction is employed in the NP-hardness proof of the problem considered in [Ahmed *et al.*(2019)Ahmed, Baali, within the nonoverlapping setting. The reduction works for our setting as well, since the reduction makes sure the output of the overlapping driver module identification in cancer problem produces a single module of size $K$. □

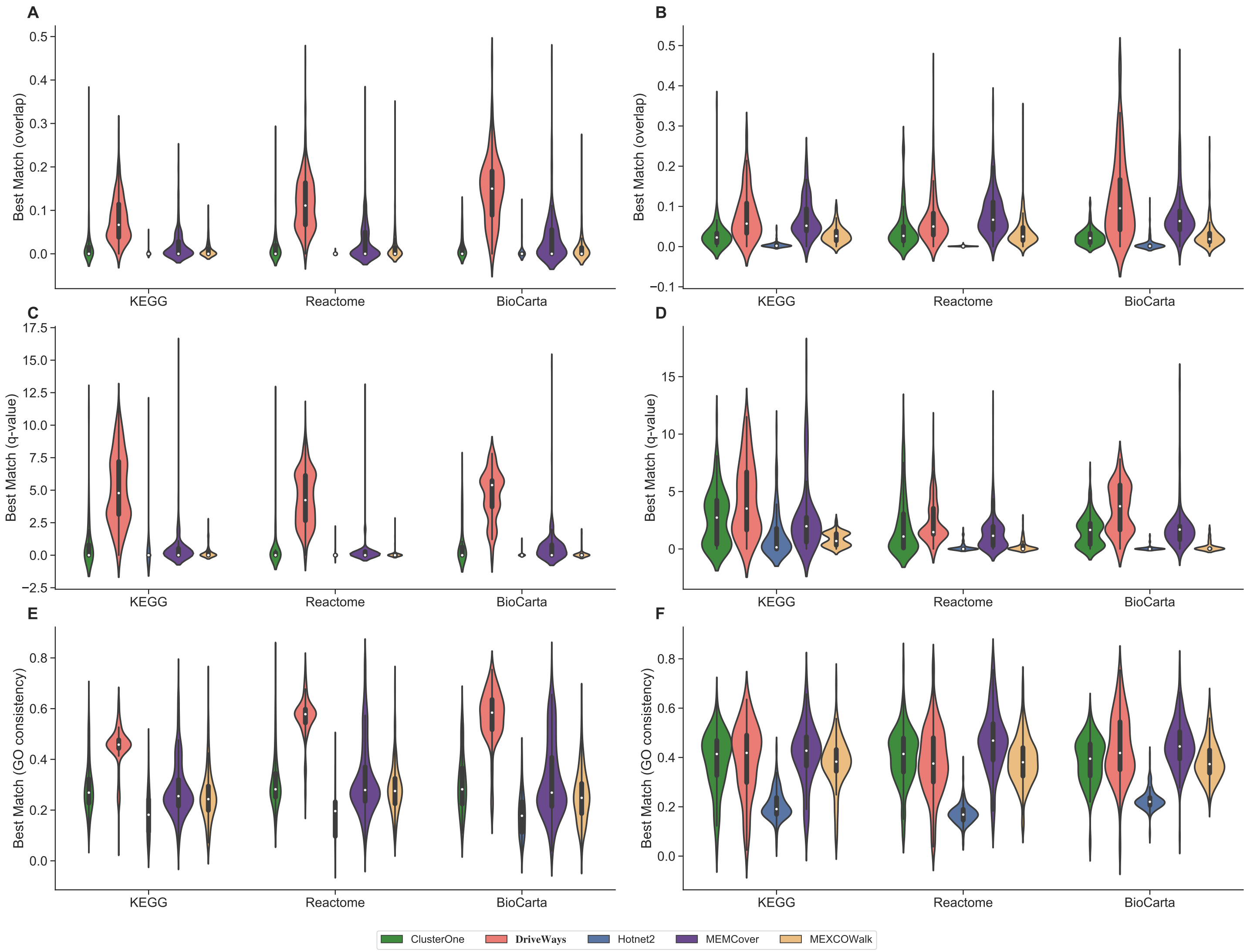


Figure S1: Evaluation of precision by comparing distribution of best match scores for each predicted module when pan-cancer samples are used as input; scores are calculated with (A) overlap score (C) q-value and (E) GO consistency. Evaluation of recall by comparing distribution of best match scores for each reference pathway when pan-cancer samples are used as input; scores are calculated with (B) overlap score (D) q-value (F) GO consistency.

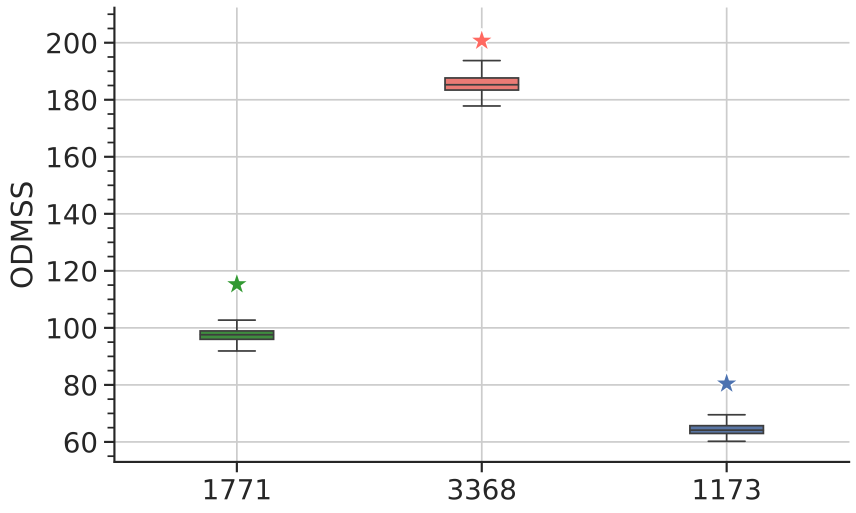


Figure S2: ODMSS results when DriveWays is run with a random seed list. ⋆ represents the results when the original seed list is used in DriveWays runs.

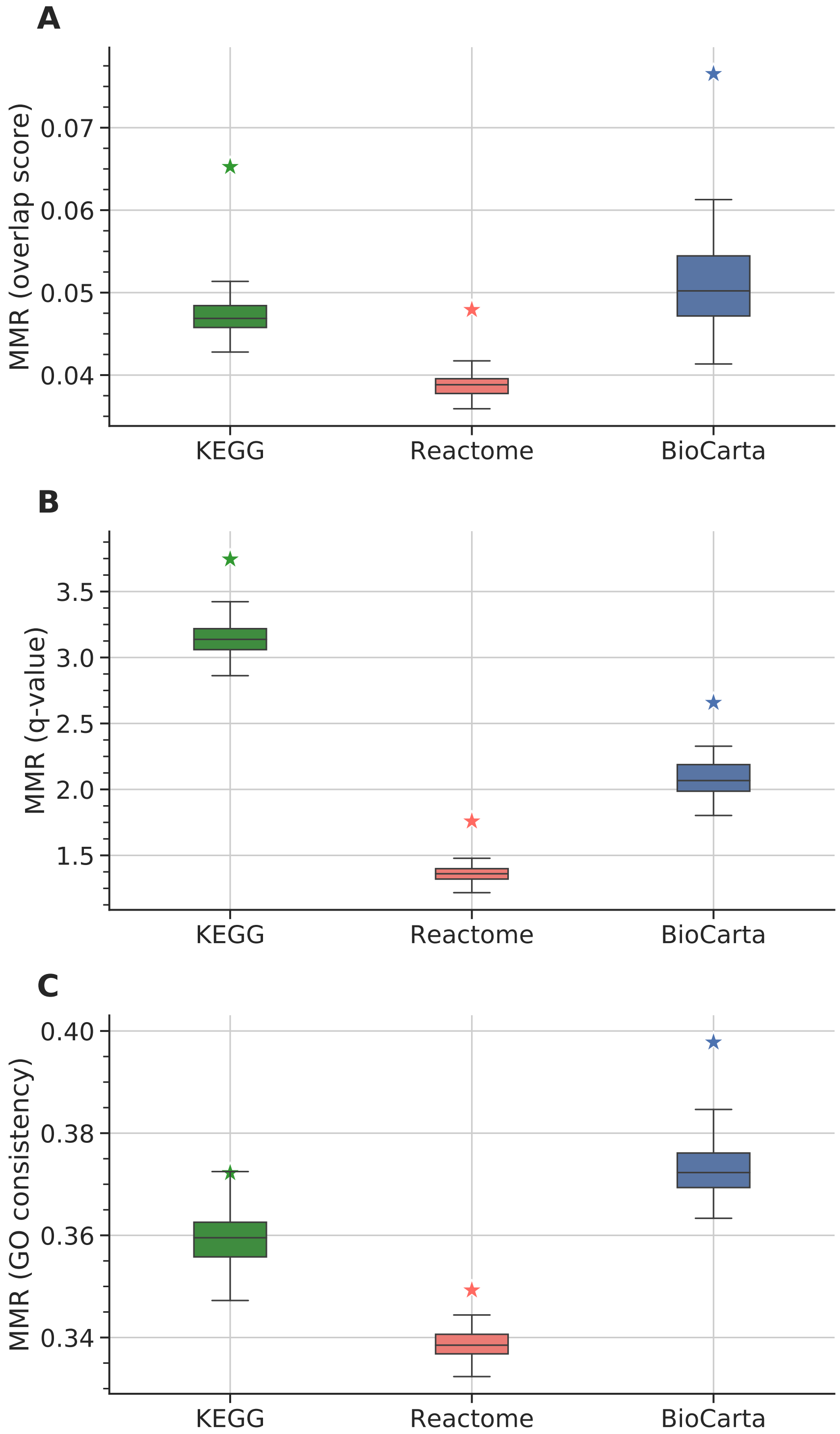


Figure S3: MMR scores when DriveWays is run with a random seed list. A) Overlap score B) Hypergeometric test q-values C) GO consistency. ⋆ represents the scores when the original seed list is used in DriveWays runs.

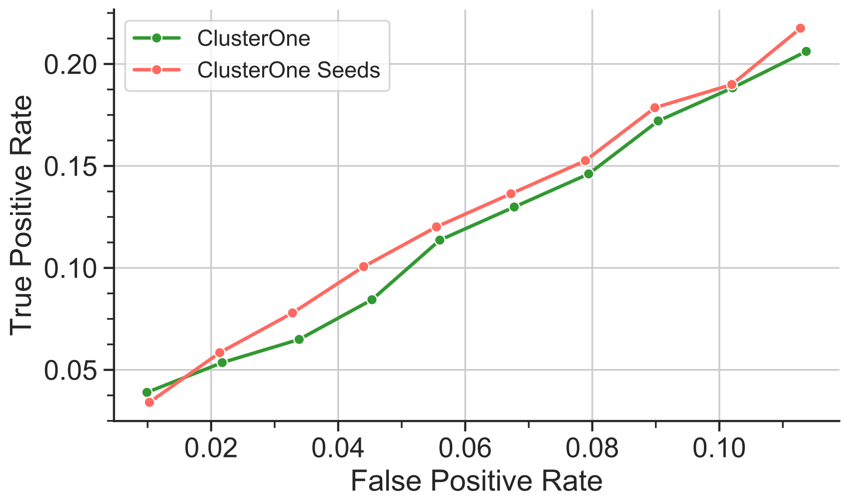


Figure S4: CGC overlap evaluation of ClusterOne output modules and ClusterOne seeds when pan-cancer samples are used as input. ROC curves are calculated for $unique\_genes = 100, 200, ..., 1000$. Note that ClusterOne seeds refer to using seeds directly without any module growth process.

#### 1.2 Results on the BRCA dataset

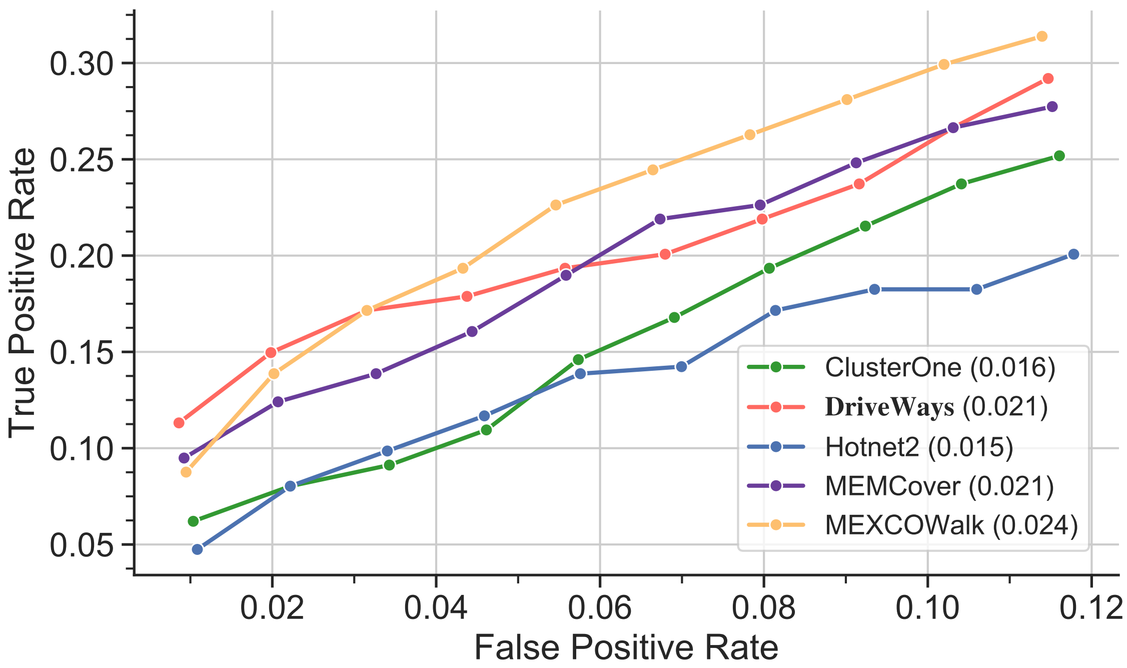


Figure S5: ROC curves calculated for $unique\_genes = 100, 200, ..., 1000$ from the outputs of considered methods when breast cancer samples are used as input and CM3 is used for reference.

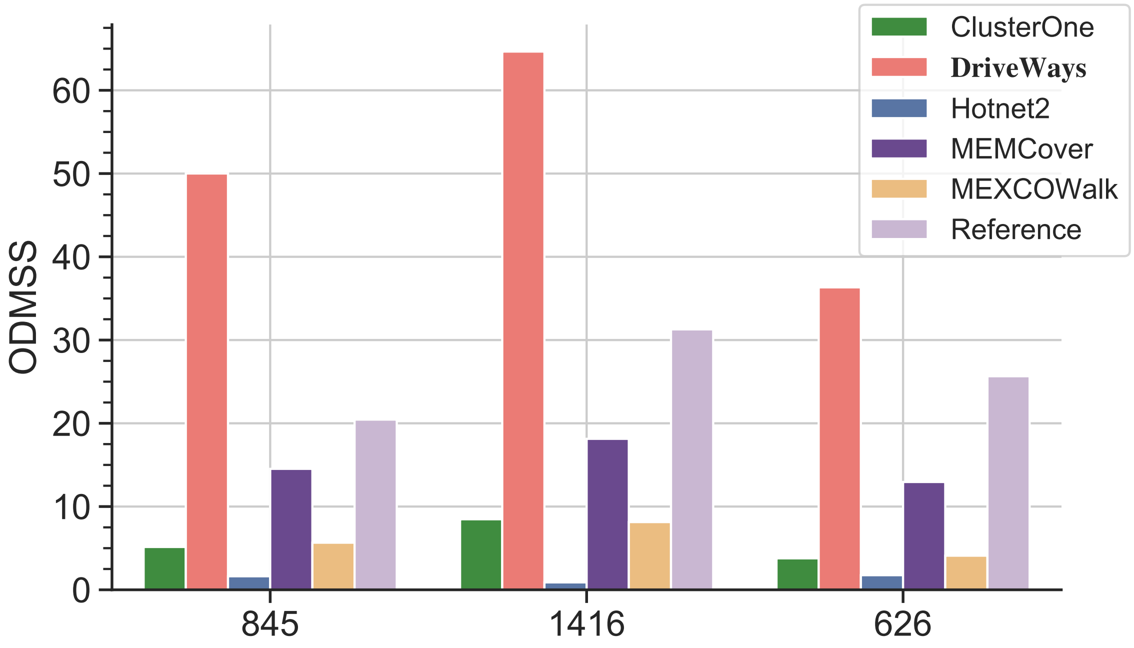


Figure S6: Overlapping driver module set score (ODMSS) obtained from breast cancer samples when $\delta_s$ parameter is determined based on the total size of $\text{KEGG}_{CM3}$, $\text{Reactome}_{CM3}$ and $\text{BioCarta}_{CM3}$ pathways, respectively.

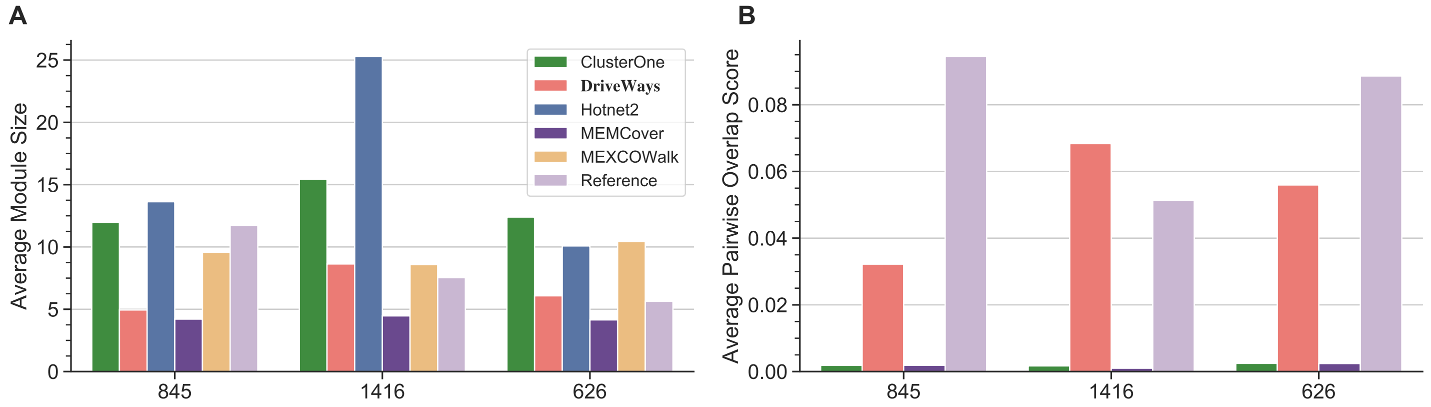


Figure S7: A) Average module sizes in the outputs of the methods under consideration for the shown $\delta_s$ values obtained from breast cancer samples. B) Corresponding average pairwise overlap scores.

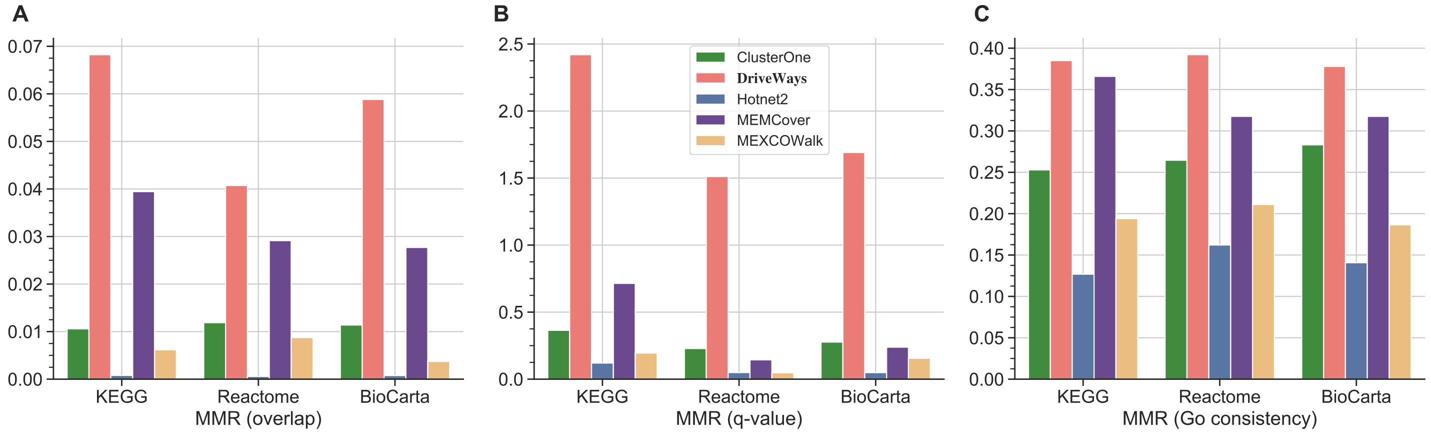


Figure S8: MMR scores of all methods calculated with three similarity metrics: A) Overlap score B) Hypergeometric test q-values C) GO consistency. The set of reference pathways at the x-coordinate of each plot correspond to $\text{KEGG}_{CM3}$, $\text{Reactome}_{CM3}$, and $\text{Biocarta}_{CM3}$, from left to right.

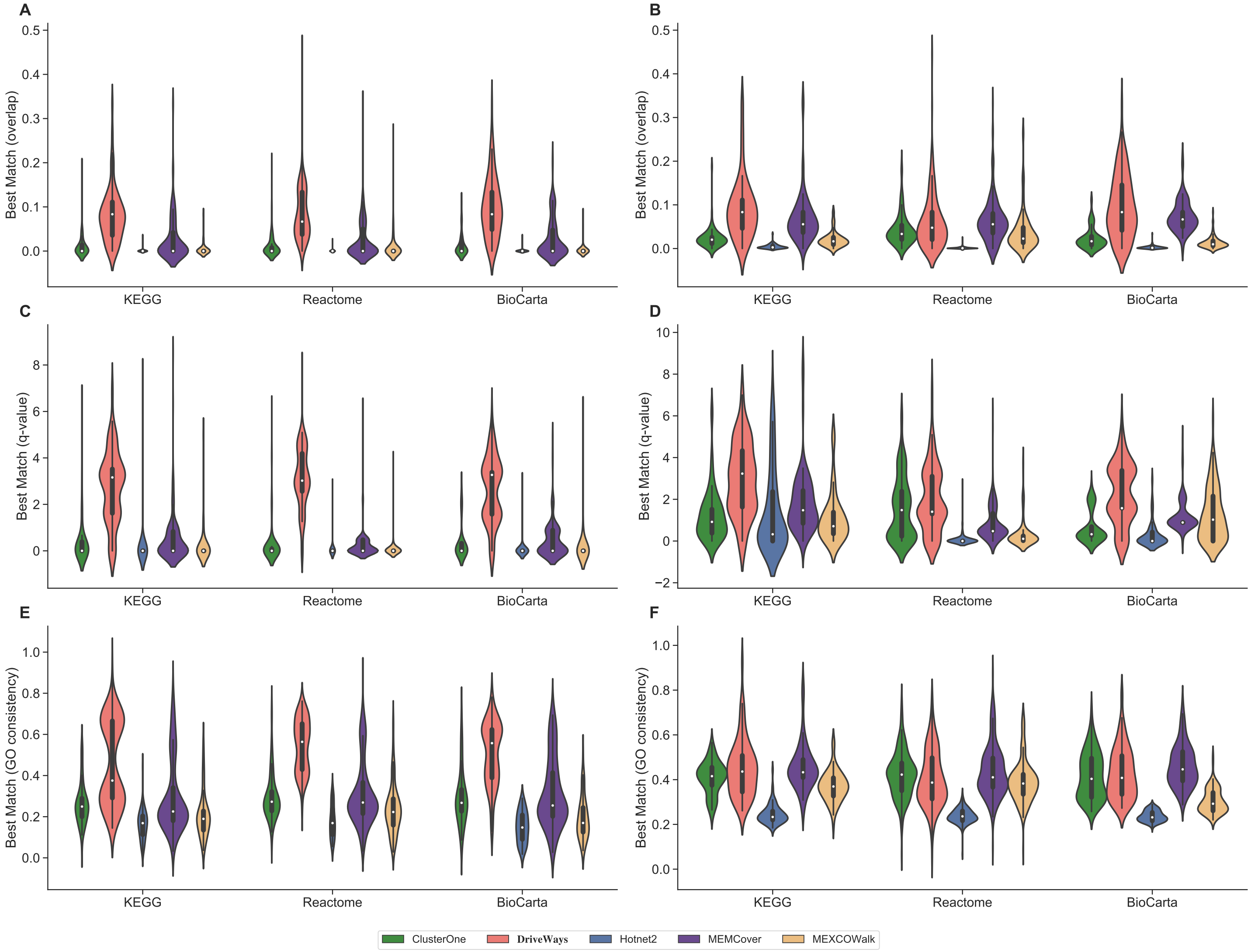


Figure S9: Evaluation of precision by comparing distribution of best match scores for each predicted module when breast cancer samples are used as input; scores are calculated with (A) overlap score (C) q-value and (E) GO consistency. Evaluation of recall by comparing distribution of best match scores for each reference pathway when breast cancer samples are used as input; scores are calculated with (B) overlap score (D) q-value (F) GO consistency.

#### 1.3 MMR Examples

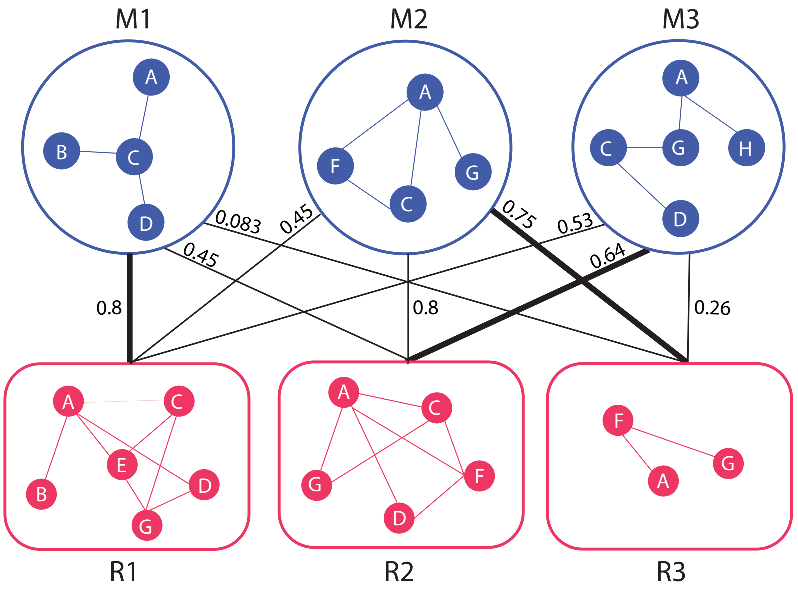


Figure S10: A toy example illustrating the calculation of Maximum Matching Ratio (MMR). M1,M2 and M3 are the predicted modules, and R1, R2 and R3 are the reference pathways. Edge weights correspond to the overlap score between the adjacent predicted module and reference pathway. Bold edges represent the edges in the maximum matching with maximum cardinality. MMR is $(0.8 + 0.64 + 0.75)/3 = 0.73$

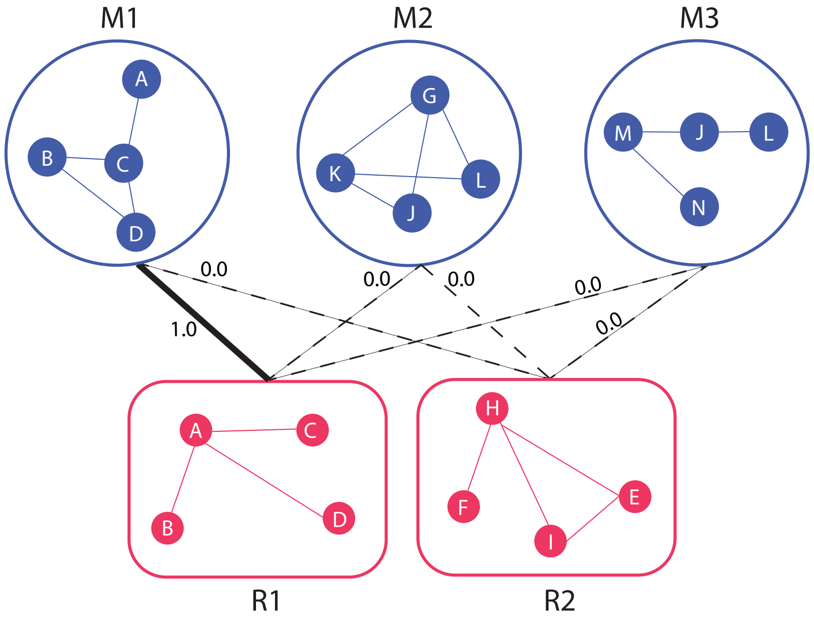


Figure S11: A toy example illustrating the effect of including zero weighted edges in the bipartite graph. If the zero edges were not present MMR would be 1, but considering those edges MMR is 1/3.

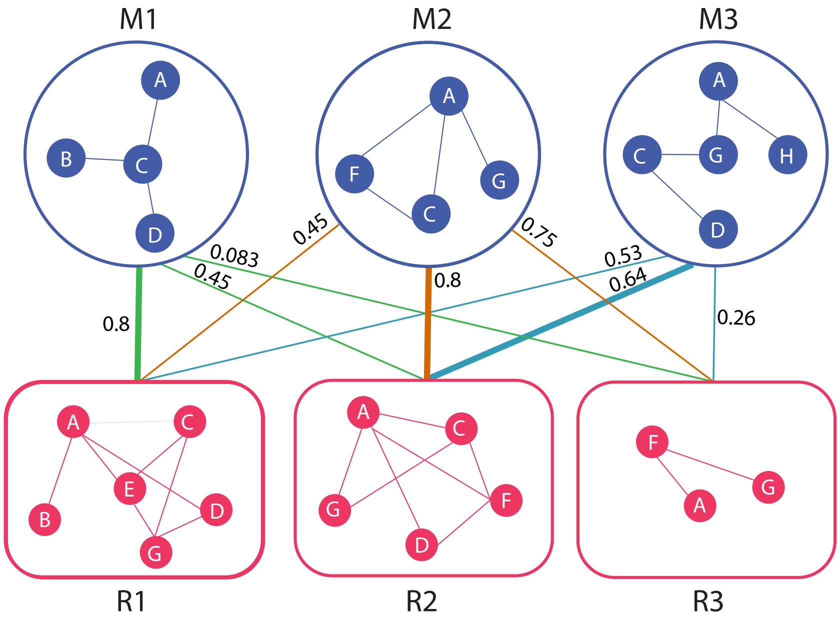


Figure S12: Illustration of finding the best match for each predicted module to evaluate precision. For each predicted module $M_i$, the best match is the maximum weighted edge between $M_i$ and $R_j$ such that $\exists j \in (1, 2, ..m)$ where $m$ is the number of reference pathways.

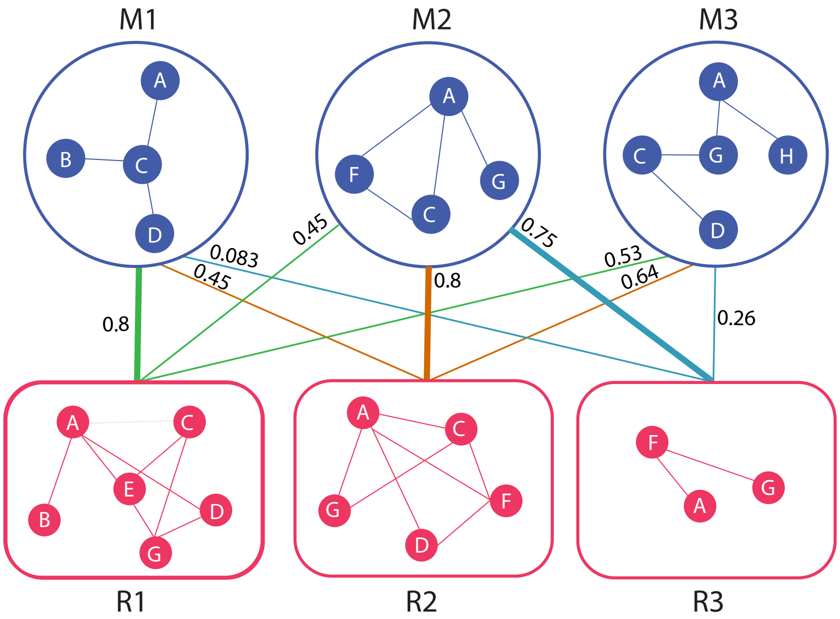


Figure S13: Illustration of finding the best match for each reference pathway to evaluate recall. For each reference pathway $R_j$, the best match is the maximum weighted edge between $R_j$ and all $M_i$ such that $\exists i \in (1, 2, ..n)$ where $n$ is the number of predicted modules.

Table S1: Statistics of reference pathway databases

| Reference | # of total genes | # of pathways | # of unique genes |
|---|---|---|---|
| $KEGG_{CGC}$ | 1771 | 104 | 330 |
| $Reactome_{CGC}$ | 3368 | 337 | 360 |
| $BioCarta_{CGC}$ | 1173 | 149 | 188 |

Table S2: Statistics on predicted modules of considered methods

| Reference | | ClusterOne | DriveWays | Hotnet2 | MEMCover | MEXCOWalk |
|---|---|---|---|---|---|---|
| $KEGG_{CGC}$ | # of modules | 118 | 327 | 40 | 395 | 216 |
| | # of unique genes | 1291 | 571 | 1770 | 1386 | 1770 |
| $Reactome_{CGC}$ | # of modules | 261 | 525 | 13 | 727 | 350 |
| | # of unique genes | 2212 | 909 | 3368 | 2649 | 3368 |
| $BioCarta_{CGC}$ | # of modules | 73 | 213 | 57 | 262 | 141 |
| | # of unique genes | 893 | 326 | 1173 | 921 | 1173 |

Table S3: Pan-cancer average F1 scores

| Reference | | ClusterOne | **DriveWays** | Hotnet2 | MEMCover | MEXCOWalk |
|---|---|---|---|---|---|---|
| $KEGG_{CGC}$ | overlap | 0.018 | **0.078** | 0.003 | 0.031 | 0.014 |
| | q-value | 1.412 | **4.589** | 0.499 | 0.856 | 0.301 |
| | GO consistency | 0.329 | **0.419** | 0.196 | 0.333 | 0.304 |
| $Reactome_{CGC}$ | overlap | 0.025 | **0.084** | 0.001 | 0.042 | 0.019 |
| | q-value | 0.915 | **3.125** | 0.156 | 0.439 | 0.108 |
| | GO consistency | 0.351 | **0.458** | 0.179 | 0.378 | 0.326 |
| $BioCarta_{CGC}$ | overlap | 0.019 | **0.132** | 0.004 | 0.048 | 0.018 |
| | q-value | 1.089 | **3.91** | 0.035 | 0.917 | 0.134 |
| | GO consistency | 0.341 | **0.495** | 0.200 | 0.374 | 0.307 |

Table S4: BRCA average F1 scores

| Reference | | ClusterOne | **DriveWays** | Hotnet2 | MEMCover | MEXCOWalk |
|---|---|---|---|---|---|---|
| $KEGG_{CM3}$ | overlap | 0.015 | **0.086** | 0.001 | 0.039 | 0.008 |
| | q-value | 0.633 | **2.848** | 0.231 | 0.766 | 0.382 |
| | GO consistency | 0.318 | **0.455** | 0.195 | 0.357 | 0.265 |
| $Reactome_{CM3}$ | overlap | 0.019 | **0.070** | 0.001 | 0.036 | 0.014 |
| | q-value | 0.491 | **2.417** | 0.060 | 0.348 | 0.116 |
| | GO consistency | 0.346 | **0.463** | 0.203 | 0.361 | 0.296 |
| $BioCarta_{CM3}$ | overlap | 0.015 | **0.097** | 0.001 | 0.041 | 0.006 |
| | q-value | 0.434 | **2.396** | 0.086 | 0.670 | 0.350 |
| | GO consistency | 0.342 | **0.463** | 0.183 | 0.373 | 0.236 |

Table S5: $t$ and $d$ values selected by BayesOpt procedure

| Reference | Input Data | $t$ | $d$ |
|---|---|---|---|
| $\text{KEGG}_{CGC}$ | | 1.043 | 2.435 |
| $\text{Reactome}_{CGC}$ | Pan-cancer | 1.0 | 3.0 |
| $\text{BioCarta}_{CGC}$ | | 1.017 | 3.011 |
| $\text{KEGG}_{CM3}$ | | 1.080 | 5.0 |
| $\text{Reactome}_{CM3}$ | Breast cancer | 0.926 | 5.0 |
| $\text{BioCarta}_{CM3}$ | | 0.986 | 3.126 |

### References


[Ahmed *et al.*(2019)Ahmed, Baali, Erten, Hoxha, and Kazan] Ahmed, R., Baali, I., Erten, C., Hoxha, E., and Kazan, H. (2019). MEXCOwalk: Mutual Exclusion and Coverage Based Random Walk to Identify Cancer Modules. *Bioinformatics*.

[Garey and Johnson(1979)Garey and Johnson] Garey, M. R. and Johnson, D. S. (1979). *Computers and Intractability: A Guide to the Theory of NP-Completeness*. W. H. Freeman & Co., New York, NY, USA.
